# The impacts of body mass on immune cell concentrations in birds

**DOI:** 10.1101/2020.04.23.057794

**Authors:** Emily Cornelius Ruhs, Lynn B. Martin, Cynthia J. Downs

## Abstract

Body mass affects many biological traits, but its impacts on immune defenses are fairly unknown. Recent research on mammals found that neutrophil concentrations scaled hypermetrically with body mass, a result not predicted by any existing theory. Although this mammalian model might predict how leukocyte concentrations scale with body mass in other vertebrates, vertebrate classes are distinct in many ways that might affect their current and historic interactions with parasites and hence the evolution of their immune systems. Subsequently, here, we asked which existing scaling hypothesis best-predicted relationships between body mass and lymphocyte, eosinophil, and heterophil concentrations—the avian functional equivalent of neutrophils—among >100 species of birds. We then examined the predictive power of body mass relative to life-history variation, as an extensive literature indicates that the scheduling of key life events has influenced immune system variation among species. Finally, we ask whether these scaling patterns differ from the patterns we observed in mammals. We found that an intercept-only model best-explained lymphocyte and eosinophil concentrations among birds; body mass minimally influenced these two cell types. For heterophils, however, body mass explained over 30% of the variation in concentrations among species, much more than life-history variation (~8%). As with mammalian neutrophils, avian heterophils scaled hypermetrically (*b*=0.19 ± 0.05), but significantly steeper than mammals (~1.5x). As such, we discuss why birds might require more broadly-protective cells compared to mammals of the same body size. Body mass appears to have strong influences on the architecture of immune systems, which could impact host-parasite coevolution and even zoonotic disease risk for humans.

## INTRODUCTION

Body size influences almost all of the behaviors, physiological processes and hence eco-evolutionary roles of species in communities [1, 2]. Nevertheless, despite the many life-history traits that we know to scale with body size and that might influence immune cell concentrations, we lack information on how body mass affects defense [3–5]. Many physiological factors that go hand-in-hand with body size evolution also likely impact exposure to parasites and pathogens and have implications for host immune systems [6, 7]. A growing body of theory and a few data propose that body size influences the immune systems of species, partly because risk of infection varies with body size [6, 7, 8]. In general, higher risk in large species occurs because large species cover greater physical distances with each movement (i.e., per step or beat of wings) and have larger respiratory, digestive and sexual tissue surface areas for infection relative to parasites [4, 8]. Large species also have distinct life-history traits from small ones (e.g. social groups, home range, feeding) [9, 10], as the only way to reach large size is via long development and subsequently delayed maturation, both of which necessitate a long lifespan [11]. Several studies have revealed that such long-living (“slow”) species (i.e. elephant; low reproductive rate and long development) have different immune defenses than short-living (“fast”) ones (i.e. mouse; high reproductive rate and short development)[12–15], such that an individual should invest in specific immune defenses if they live for long periods because they may come into contact with the same pathogens repeatedly. Yet to what degree these life-history patterns are genuine or just echoes of body mass effects remains unclear.

In the present study we investigated whether and how body mass is related to immune cell concentrations among birds spanning several orders of magnitude in mass (~1180 fold; Fig. S1). Our particular interest was to identify which hypothesis best predicted the scaling relationships for immune cell concentrations among birds (Fig. 1), and then to test whether these scaling patterns are different from those found in mammals [16]. The main parameter of interest is *b*, or the slope of the line relating to body mass and leukocyte concentration. For many traits, scaling relationships are isometric, such that changes in traits are directly proportional to changes in body mass. Other traits, though, scale hyper- or hypometrically, such that the trait of interest is disproportionally larger or smaller than would be expected by geometry. Two hypotheses predict isometric scaling for immune cell concentrations (*b* = 0, black dashed line in Fig. 1). The first is the Protecton hypothesis, which proposes that all organisms require similar levels of protection, regardless of size [17, 18]. Similarly, the related Complexity hypothesis [16] assumes that the time for surveillance of tissues and delivery of a single leukocyte is independent of body mass [19, 20]. Another hypothesis, Rate of Metabolism [21], invokes metabolic rate as the key driver of variation in immune cell concentrations [1]. Thus, the rates of proliferation and actions of immune cells should be reliant on the Rate of Metabolism of a host [21]. This hypothesis predicts hypometric relationships between leukocyte concentration and body mass (*b* = −0.25, red line in Fig. 1). A final hypothesis is derived from our recent discovery about neutrophil scaling in over 250 species of mammals [16] and prior theory on the evolution of similar organismal functions [22]. In our previous study [16], we found that neutrophil concentrations scaled hypermetrically with body mass (b=0.11; blue line in Fig. 1). This particular outcome led us to offer the Safety factor hypothesis [23], which proposes that hypermetric scaling may have evolved, in particular for early anti-microbial defenses, because larger species require disproportionately more rapidly-acting, broadly-protective defensive cells to combat infections than small species [24]. In other words, we expect that this pattern evolved because large species must prioritize risk reduction over other life-history priorities because they mature more slowly and hence live longer than smaller species [11, 12, 16].

**Figure 1.**
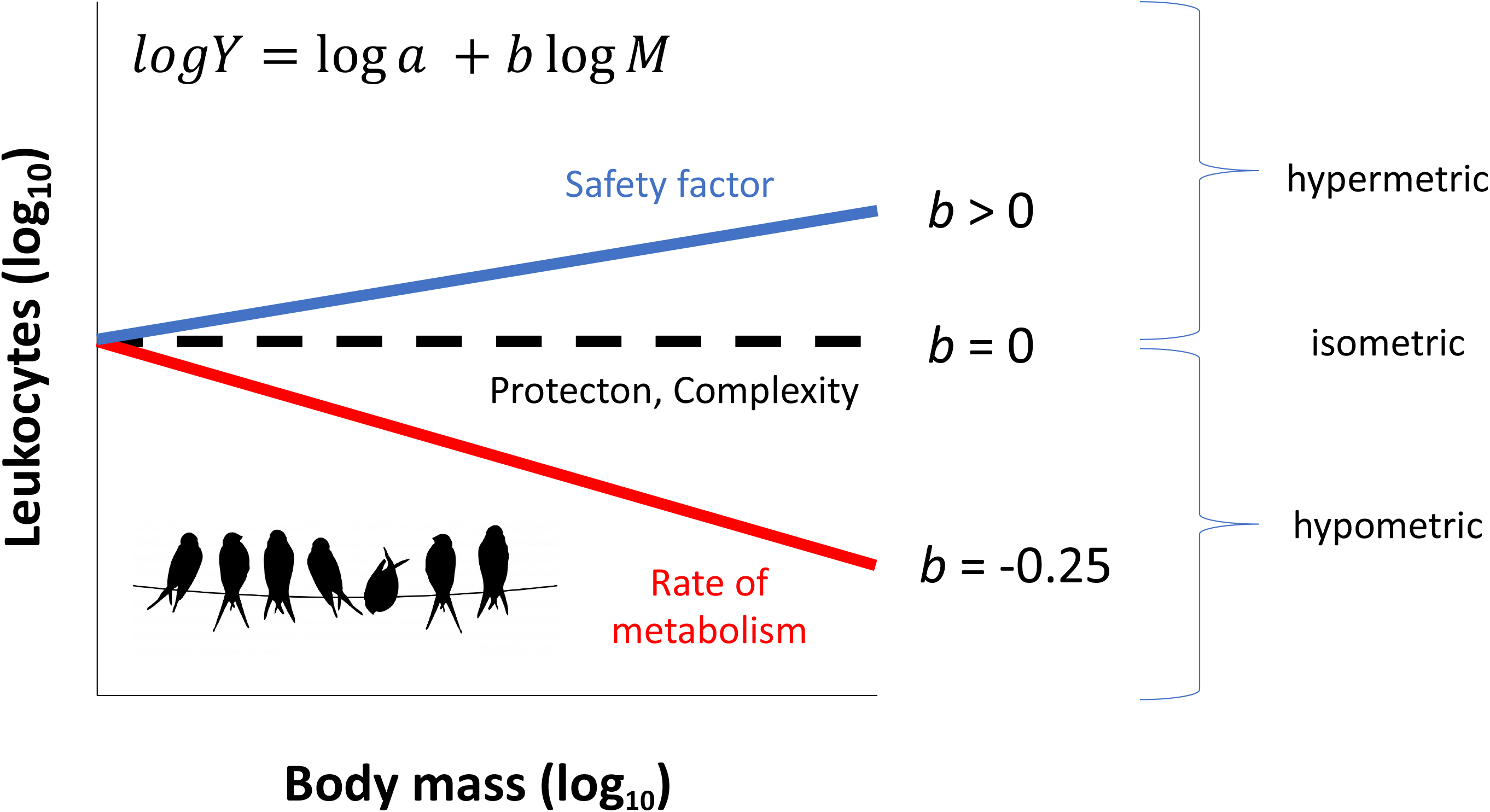
Possible scaling relationships between immune cell concentrations and body mass. Classic work from Kleiber (1932), Calder (1981), Schmidt-Nielson (1984) and others has shown that many biological traits vary consistently with body size [5–7]. Linearized forms of the equation, *Y* = *aM^b^*, are typically used to describe such relationships where *a* represents the intercept, *M* connotes body mass, and *b* represents the scaling exponent. The equation in the figure represents these linearized functions with each curve on the figure highlighting the main parameter of interest in the present study, *b*, or the slope of the line relating body mass and leukocyte concentrations. For rate processes and concentrations, *b* estimates tend to be negative or in other words, they scale hypometrically (*b* < 0; red line). Perhaps one of best-known hypometric relationships is that of body size and metabolic rate (b=-0.25)[5]. For other traits, scaling relationships are isometric (*b* = 0; black dashed line), such that changes in traits are directly proportional to changes in body mass, or much more rarely, scaling relationships are hypermetric (*b* > 0; blue line). In the present study, we sought to determine how immune cell concentrations relate to body size among birds. Hypotheses for immune defense scaling in birds (*b* exponents in figure) were based on theory or patterns observed in other taxa. The three slope estimates depicted represent distinct hypotheses (Safety factor, Protecton/Complexity and Rate of Metabolism), which are described in detail in the text. Inset bird image from https://pixabay.com/illustrations/bird-birds-wire-perched-perching-220327/.

Here, we sought to determine which hypothesis best predicted scaling relationships for three leukocyte concentrations (heterophils, eosinophils, and lymphocytes) among avian species. To enable comparison with published values describing how immune cells scale with body mass in mammals, we used the same approach as Downs et al. (2020). To our knowledge, there is only one other study that has investigated this relationship across various species of birds; however, these data are gathered from wild species and not phylogenetically controlled [25]. Therefore, there is a need to understand how phylogeny and other life-history characteristics contribute to allometric patterns in similarly-housed healthy individuals. Specifically, we conducted three modeling exercises. First we asked which hypothesis best-predicted scaling relationships: the Protecton/Complexity hypothesis (isometric slope), the Rate of Metabolism hypothesis (hypometric slope), or the Safety factory hypothesis (hypermetric slope) [16]. For the Safety factor hypothesis, we did not attempt to fit the same *b* as discovered in the mammalian study for heterophils, and instead estimated the *b* from the data. We took this approach because birds and mammals differ in many ways that could influence the magnitude (or even direction) of the slope. For instance, birds do not possess neutrophils, but instead maintain a functionally similar granulocyte called a heterophil. Neutrophils are derived from vascular spaces, which overall could provide mammals a potentially bigger cell pool (outside general circulation) on which to draw granulocytes [25]. Heterophils, by contrast, are derived from stem cells in the extravascular spaces of bone marrow or by extramedullary hematopoiesis. Heterophils are fast-acting leukocytes able to engulf or control microbes, typically by mechanisms involving oxidation, upon first exposure [33, 45, 46, 48]; we thus expected similar hypermetric scaling as with mammals. Eosinophils also have diverse roles but are especially important in the control of helminth infections and other extra-cellular parasites [33]. Their functional resemblance to heterophils also led us to expect hypermetric scaling for this leukocyte type. Compared to the other two cell types, lymphocytes are exceptionally heterogeneous [33], an amalgam of quiescent and active B and T cells with diverse protective roles against intra- and extra-cellular macro- and microparasites through various mechanisms from the production of antibodies, to the coordination of other immune responses, to direct control of viral pathogens. For this leukocyte class, we expected the predictive power of body mass to be weak given the high functional diversity of this cell group. Most birds also lack the differentiated lymph nodes of mammals, and instead maintain diffuse lymphoid tissue that creates local nodules wherever antigen-stimulation occurs [27]. Birds might therefore lack a tissue repository for some leukocytes [28], which would affect concentrations of leukocytes in circulation.

Our second modeling exercise sought to discern whether body mass or life-history traits (maximal lifespan, maximal reproductive capacity, and interaction with body mass) best predicted leukocyte counts in birds. We expected body mass to predict variation in all three cell types better than life-history variables, but we expected phylogeny also to be a strong predictor as we and others have observed previously [16, 29–32]. However, we also expected predictor effects to vary among cell types based on their respective defensive functions. Our third and final modeling exercise directly tested whether the scaling patterns observed in birds and mammals for lymphocytes and heterophils/neutrophils differed from one another.

## MATERIALS AND METHODS

### Trait data

We extracted species means of heterophil and lymphocyte concentrations (cells L^−1^) from whole blood for 116 avian species and eosinophil concentrations for 88 avian species from Species360 (Table S1) [34]. Cell concentration data come from captive, adult animals housed in zoos accredited by the Association of Zoos and Aquariums (AZA) and considered healthy [34]. We used data from adult, captive individuals to minimize the confounding effects of infection and unknown reproductive status that would have arisen if we studied wild individuals. Zoo collections are biased toward large-bodied birds. Consequently, this database underrepresents small birds (for a comparison to the world’s bird species body mass distributions see Fig. S1).

We were conservative and used only Global Species Reference Intervals and absolute counts reported for each species (see Dryad for descriptions). We extracted life-history data on body mass, maximum lifespan, age at maturation, inter-laying interval (i.e., how often a species lays a clutch), average clutch size and incubation period from the CRC Handbook of Avian Masses [35] and publicly available databases such as AnAge [36], and the Animal Diversity Website [37]. Data were compiled as best as possible using a combination of all sources so as to have the most complete dataset. From these life-history data we calculated maximal reproductive capacity for each species, a similar metric to our previous study of mammals [16]:

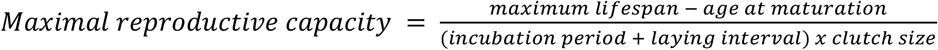

### Correlation analyses

To determine if cell concentrations were correlated, we conducted Pearson’s correlation analyses between log_10_-transformed (1) lymphocytes and heterophils, (2) lymphocytes and eosinophils and (3) heterophils and eosinophils. We also performed correlations among body mass and life-history traits.

### Exercise 1. Leukocyte allometry modelling

Our first interest was to determine which hypothesis best-predicted scaling relationships for three leukocyte concentrations among avian species. Therefore, we fit three sets of *a priori* models to the log_10_transformed concentrations of lymphocytes, heterophils and eosinophils, separately (9 total models):

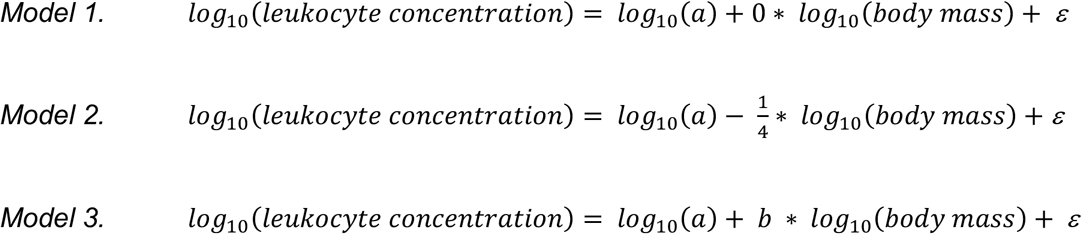

Model 1 was an intercept-only model that corresponded to the Protecton/Complexity hypothesis and hence an isometric slope (*b* = 0). It was also the model we would expect based on a null hypothesis that leukocyte concentrations scale in direct proportional to body mass. Model 2 corresponded to the Rate of Metabolism hypothesis, which predicts hypometric scaling (*b* = −0.25). Model 3 estimate the slope from the data which allowed us to test for hypermetric scaling predicted by the Safety factor hypothesis.

### Exercise 2. Comparing effects of body mass relative to life-history

To determine the ability of body mass to explain concentrations of heterophils, eosinophils, and lymphocytes relative to life-history traits, we fitted three separate omnibus models (one for each cell type) that included log_10_(body mass), maximum longevity, maximum reproductive capacity, and all two-way interactions between log_10_ (body mass) and life-history variables as fixed effects.

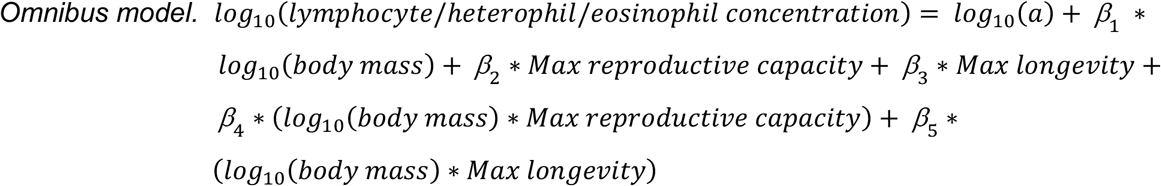

### Comparing models and determining fit

All analyses involved phylogenetic mixed effects models in R (version 3.6.0) using the packages ape [38] and MCMCglmm [39]. For modelling exercises 1 and 2, we included phylogenetic effects from a tree we produced by pruning the time-rooted phylogenetic tree created by Uyeda et al. [40] to our species list, then using that tree to create a phylogenetic covariance matrix. Our sample sizes differed among cell types (n=116 for lymphocytes and heterophils and n=88 for eosinophils), due to differences in availability and completeness of data in Species 360 and our life-history references. Therefore, we created and used two different phylogenetic trees, one for lymphocytes and heterophils and another for eosinophils (Fig. S2). We set the inverse-gamma priors to 0.01 for the random effect of phylogenetic variance and default priors for the fixed effects in all models except Model 2, where we used *a priori* value of −0.25 [21]. Models were run for 260k iterations with 60k burn-in and a 200-iteration thinning interval.

We calculated model fits for all models from exercise 1 and 2 using conditional R^2^ [41], and then used Deviance Information Criterion (ΔDIC) to identify the best-fit model. We first compared models 1-3, we defined the top model as the model with the lowest DIC and considered models within a ΔDIC<2 as having equivalent support. We then compared the top model from exercise 1 with the omnibus model for each cell type. This approach enabled us to discern whether previous discoveries for life-history variable effects on immune variation might mostly be due to body mass. For all models, we also calculated (1) Pagel’s lambda to determine the amount of variation explained by the phylogeny accounting for fixed effects, (2) Pagel’s unadjusted lambda to determine the variation explained by the phylogeny not accounting for fixed effects, and (3) marginal R^2^ values to determine how much variation in leukocyte concentrations was explained by fixed effects [42; 41]

### Exercise 3. Direct comparison of bird and mammal slopes

Our third modeling exercise was designed to compare the slopes of the relationship between body mass and cell type in birds (n=116) and mammals (n=259). The mammal dataset included only lymphocytes and neutrophils, limiting our analyses to those cell types. We compared heterophils and neutrophils because they have similar functions [43]. We created a new phylogenetic covariance matrix estimated using a phylogenetic tree constructed with data from the National Center for Biotechnology Information and phyloT [44]. Polytomies were resolved during tree construction using the randomized function built into phyloT. The bird/mammal model was fit with an inverse-gamma prior set to 0.01 and run with burn-in, iterations and a iteration thinning interval similar to Model 3. However, we included the fixed effect term Class and the interaction between log_10_(body mass) and class. For the two models (lymphocytes and hetero-/neutrophils, separately) we then calculated Pagel’s lambda, Pagel’s unadjusted lambda, and marginal R^2^ values.

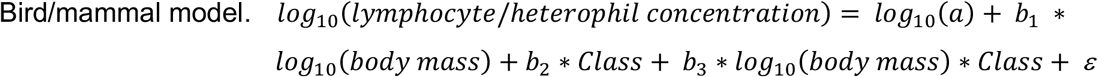

## RESULTS

### Correlation analyses

Log_10_-transformed lymphocytes did not covary with log_10_-transformed heterophils (r = 0.102, t_117_ = 1.11, p = 0.269), nor with log_10_-transformed eosinophils (r = 0.164, t_86_ = 1.54, p = 0.127). However, species with more log_10_-transformed heterophils had more log_10_-transformed eosinophils (r = 0.402, t_86_ = 4.07, p = 0.0001). We also found correlations between log_10_-transformed body mass and life-history traits (Fig. S3).

### Best-fit models for leukocyte allometry

The Protecton/Complexity model (Model 1) best-predicted avian lymphocyte and eosinophil concentrations (Table 1), but the Rate of Metabolism model (Model 2) was within 2 ΔDIC for both leukocyte types (ΔDIC=-1.25 for lymphocytes and −0.97 for eosinophils) [4]. However, body mass was a weak predictor of variation in lymphocytes and eosinophils (Model 3; Fig. S4), because it explained <0.1% much variation in either lymphocyte or eosinophil concentrations. Overall, Model 1 (fitting *b=0*) accounted for 69% and 91% of variation in lymphocytes and eosinophils, respectively (Table 1).

**Table 1.**
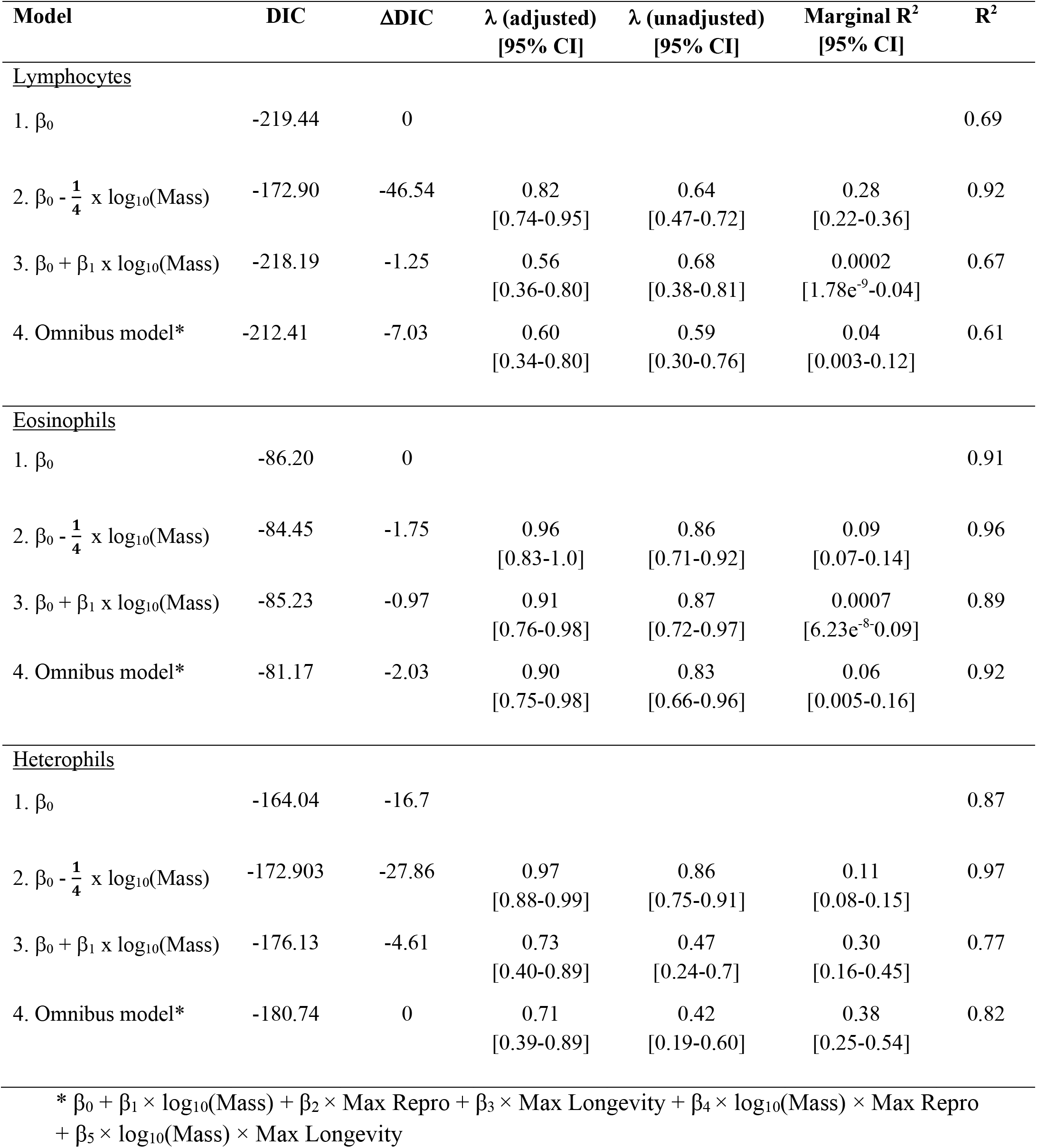
Best-fit models predicting circulating leukocyte concentrations in songbirds. Top models are represented by 0 ≤ ΔDIC ≥ 2. Model 1 fit a slope of 0, model 2 a slope of −0.25 and models 3 and 4 were allowed to estimate the slope given the data. Models test for the effects of body mass and life-history metrics on log_10_-transformed lymphocyte, eosinophil and heterophil concentrations.

Outcomes differed for heterophil concentrations (Table 1, Fig. 2A). First, the inclusion of body mass improved model fit, when the slope coefficient was estimated from the data (Model 3). Model 3 accounted for 77% of the variation in heterophils, with body mass alone accounting for 30% of that variation. In this model (but also in the omnibus model; see below), heterophil concentrations scaled hypermetrically (Model 3: *b*, 95% credible interval = 0.19, 0.14-0.24; Fig. 2A). In all models for all leukocyte types, appreciable phylogenetic effects were detected (Table 1).

**Figure 2.**
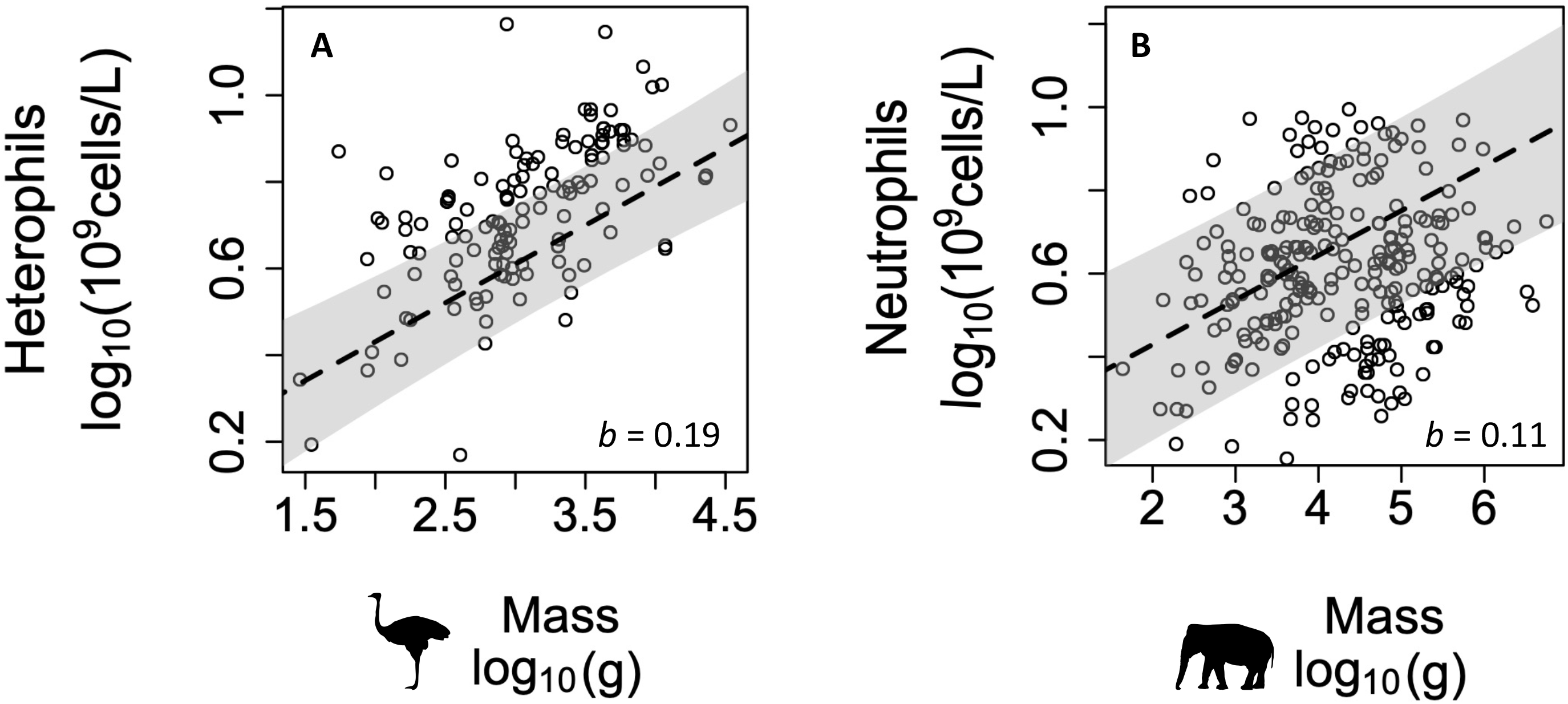
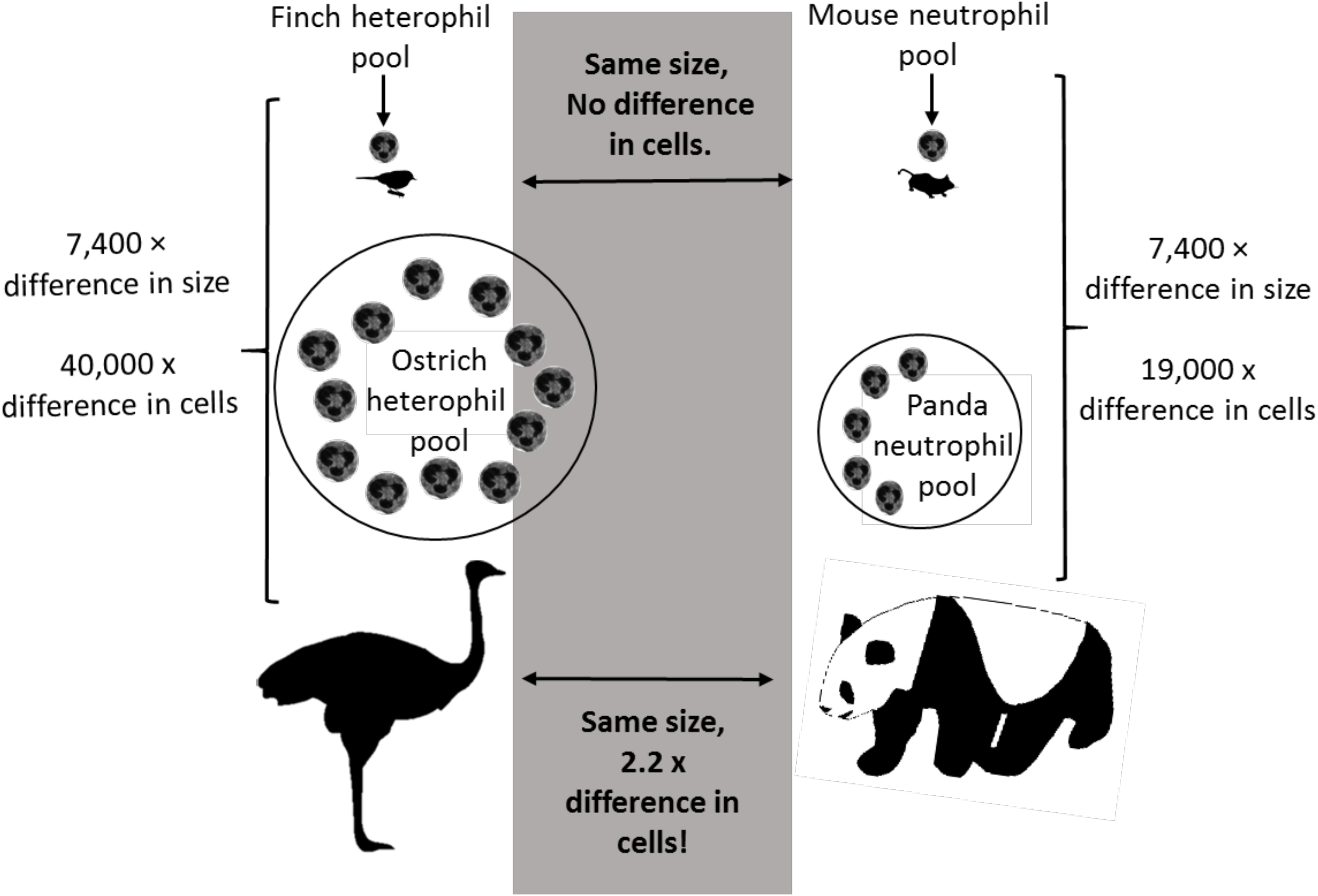
Observed scaling relationships between body mass and (A) heterophils from birds and (B) neutrophils from mammals, and (C) a visual summary of heterophil/neutrophil concentration scaling differences between birds and mammals. Shaded areas in A and B depict 95% credible intervals of the slope estimates from model 2 (mass-only). The shaded comparisons in C highlight that the heterophil/neutrophil concentrations in a finch and mouse are predicted to be similar, but an ostrich circulates 2.2-fold more cells than a giant panda.

### Effects of body mass relative to life-history

The lymphocyte and eosinophil omnibus models (model with life-history variables as fixed effects) were not amongst the top models (Table 1; ΔDIC=-7.03 and −2.03, respectively). The heterophil omnibus model was the overall best-fit (Table S2), accounting for 82% of variation in heterophils (Table 1). However, the addition of life-history traits increased explanatory power of fixed effects by only 8% (Table 1). In light of these subtle effects, we selected Model 3 (mass-only; estimating *b* from the data) as the more informative and simpler alternative about the scaling of heterophils.

### Direct comparison of bird and mammal slopes

The bird/mammal model for lymphocytes accounted for 80% of the variation (Table S3); however, the fixed effects, body mass and class, explained only 2% of the variation (Table S3). The model for hetero-/neutrophils explained 92% of the overall variation (Table S3) with body mass and class explaining a higher proportion of the variation (11%). Hetero-/neutrophils scaled hypermetrically with body mass (*b*, 95% credible interval = 0.19, 0.14-0.24) and the interaction between body mass and class predicted cell concentrations (Table S4). Taken together, these results suggest that the scaling coefficient for the relationship between body mass and hetero-/neutrophil concentrations is steeper in birds (*b*=0.19) than in mammals (*b*=0.11) [16].

## DISCUSSION

Here, we investigated the scaling of three types of leukocytes across 116 species of birds. While our database underrepresents small birds due to biases in zoo collections, it does represent 21 orders and a 1180-fold difference in size. We found little evidence for allometry for lymphocyte and eosinophil concentrations; the best-fit models for both cell types did not include body mass. These results are consistent with the Protecton hypothesis, but we encourage caution in interpreting these results as support for it. It is difficult to use null models as evidence of isometry as it is unclear whether the slope is truly zero (isometry) or if there is an absence of a pattern. Heterophils, by contrast, scaled hypermetrically, although the omnibus model performed better and explained more of the variation (82%) than the model with only body mass as a fixed effect (Model 3). These results indicate that body mass explains much more variation in heterophil concentrations (~30%), compared to life-history traits (~8%). Our most striking discovery was the slope of the effects of body mass on heterophil concentration among birds (*b*, 95% CI = 0.19, 0.14:0.24). This coefficient supports the Safety factor hypothesis, but it was also significantly steeper than the estimate we described previously for mammals (*b* = 0.11; Fig. 2B). Below we discuss the ramifications of these results, in particular why avian heterophils scale so much more steeply than mammalian neutrophils. This difference between mammals and birds was also supported by the direction comparison made in this study.

### Lymphocyte and eosinophil scaling

Lymphocytes are a heterogenous class of leukocytes, including both B and T cells. In fact, in broiler chickens, in peripheral blood, 45.9% are T-cells and 39.2% are B cells [45]. These cells have such diverse responsibilities and can be in such different states of activation that it might be difficult to detect relationships between body mass and lymphocyte concentrations even if they exist. However, allometries have been detected for lymphocytes in several but not all other studies (all mammals (*b=-0.07* [46]; *-0.04* [16], primates (*b=-0.1* [19, 32]), carnivores [47], birds [25] and bats [48], but not rodents [49]). Avian and mammalian lymphocytes provide predominantly the same forms of protection [50], so it is interesting that such different patterns of scaling have been detected among species. However, these scaling exponents might misrepresent biological relevance in the sense that many of them explain little of the variation and some do not account for the variance explained by phylogeny. For instance, in our previous mammalian study, body mass explained only 3% of variation among species in lymphocyte number [16] and the only other large scale comparison of birds does not use phylogenetically-informed statistics [25]. It would be quite valuable to develop tools to characterize counts of classes of lymphocytes that can be applied to phylogenetically-broad species that could enable us to study scaling of individual lymphocyte classes, like B and T cells.

The above logic for lymphocytes probably does not transfer to eosinophils. An absence of scaling for eosinophils is less likely derived from functional diversity among members of this class because all eosinophils perform the same type of defense. Eosinophils respond rapidly to local inflammation, mostly when it is induced by macroparasites to which their defensive actions are best attuned [51]. In birds, we detected no effect of body mass on eosinophil concentrations, but this finding is inconsistent with work in bats [48], carnivores [47], and primates (*b=0.05* [32]), all of which detected hypermetric scaling. The largest study to date in birds also found slight hypermetric scaling; however, the data were not log-transformed; (*b*=0.048) [25]. This discrepancy could be due to at least three factors. First, as with lymphocytes, body mass might have statistically significant but subtle influences on eosinophil concentrations in some taxa. Second, eosinophils are more common in tissues than the blood [51] (these data are from whole blood); therefore, scaling patterns might be more prominent in tissus. Third, eosinophils generally circulate in low numbers in both birds and mammals [51], and there might be simply too little variation within species to detect patterns across some clades. That is, a floor effect on total concentrations could impede estimation of *b* for this cell type.

### Heterophil scaling

Our most compelling discovery involved the hypermetric *b* for heterophils. Heterophils, the avian functional equivalent of mammalian neutrophils, are phagocytes that protect predominantly against microparasites. Much of our knowledge about heterophils comes from work on mammalian neutrophils, and most evidence indicates that heterophils and neutrophils share hematopoietic history and function [52]. Among mammal species and wild birds, neutro-/heterophil concentrations consistently scale hypermetrically, (e.g., bats [48], rodents [49], carnivores [47], primates [32], mammals broadly [16] and wild birds [25]). Previously, we argued that this pattern might have evolved because large mammals traverse greater distances and have higher surface area for infection while also experiencing disproportionate disadvantages relative to their parasites [4, 6]. Especially for bacteria and viruses, replicative and evolutionary advantages of infective adversaries might have forced large hosts to evolve especially robust, generic constitutive defenses [8, 32, 46]. Small hosts, by contrast, might circulate comparatively fewer of this cell type as a bet-hedge against such risks (given their inherently shorter lifespans), perhaps because they remain somewhat more capable of keeping pace in arms races with parasites because of their life-history strategy. Future studies should strive to test these hypotheses, but at the same time, attempts should be made to resolve directly how body mass interacts with life-history traits to influence interspecific variation in immunity. As above, life-history traits have been claimed to drive immune variation among species and populations, but almost never have body mass effects on life-history been disentangled from body mass [7]. Among the bird species studied here, we found some correlations between body mass and life-history traits (Fig. S3). To understand why hypermetric scaling exists at all, though, it will be important to study body mass and life-history traits together, as large organisms cannot become large without a long developmental period and subsequent selection for low rates of mortality [11].

### Why do avian heterophils scale so steeply with body mass?

Birds and mammals differ in many ways which might explain why heterophil concentrations in birds scale more steeply than neutrophil concentrations in mammals. For instance, birds are much better able to mitigate the costs of oxidative damage than mammals, which might be one explanation for their relatively longer lifespans for mammalian species of similar size [53; 54]. The majority of such differences between classes, however, would be expected to influence only the *intercept* terms in the models we evaluated, not the slopes. In other words, whereas similar-sized birds tend to live longer than mammals [9], if such a difference influenced leukocyte concentrations, its effects would be expected to do consistently across body masses, not disproportionately at large sizes as we observed. For most differences between birds and mammals (i.e. air sacs versus lungs for breathing, egg-laying versus live birth, organ mass, blood circulation/volume, and many more), differences in avian heterophil concentrations would be consistently different from mammalian neutrophils across all body masses. Our discovery of a fairly striking difference in the *steepness* of scaling between avian and mammalian granulocytes and mass was unexpected; *b* for birds in our mass-only model (Model 3) was ~1.5x the mammalian estimate from a comparable model.

When data are plotted on log-log scales (Fig. 2a&b), their biological significance is often obscure, so to convey better the consequence of this difference, we produced conceptual Fig. 3 using commonly known species at opposite ends of the extant bird species body mass distribution. Using the equation from Model 3 with published values for blood density and volume for birds [9, 34, 55], we first calculated total whole-animal heterophils for species. We estimated that an ostrich (*Struthio camelus;* 111000g) circulates 33.9k-fold more heterophils than an American goldfinch (*Carduelis tristis;* 15g), although their body masses differ by just 7.4k-fold. To capture the consequences of this allometric difference from what we observed previously in mammals [16], we tallied the number of cells in avian and mammalian species of the same size. We found that whereas a mouse (*Mus musculus*) and a finch would maintain roughly the same number of neutrophils/heterophils in their blood, an ostrich would circulate 2.2-fold more heterophils (1.17×10^20^ cells) than a giant panda (*Ailuropoda melanoleuca*) circulates neutrophils (5.43×10^19^ cells; Fig. 3C).

Our approach cannot reveal *why* large birds circulate so many more heterophils than large mammals circulate neutrophils, but there are a few conspicuous possibilities. The first is that the two cell types are not functionally similar after all, and for some reason, avian heterophils might require disproportionate compensation for less effective function via increases in cell number as birds increase in size. The second possibility involves mode of locomotion. As most birds fly, a bird would presumably expose itself to more risk space for infection than a similarly-sized mammal. One way to test this hypothesis would be to investigate neutrophil scaling in bats; such research has occurred, but allometric slopes were not estimated. Of course, though, the largest birds do not fly, and investigation of their position on scaling curves does not support this perspective. A third possibility entails the lack of lymph nodes and leukocyte pools in birds. Because birds lack defined lymph nodes, they might not be able to shuttle heterophils as efficiently around their bodies, particularly at large body sizes, as mammals can do for neutrophils. Until we know how lymphoid tissue is organized relative to body mass in birds, we can only speculate that it might scale more hypometrically than mammals. In mammals, we know that myelocytes (leukocyte progenitors) can reside in a “lazy pool,” and mature neutrophils can persist in a “rapid mobilizable pool”[28]. The existence of heterophil pools in birds is unknown, but as we proposed for lymphoid tissue, steep hypometric scaling of the size or numbers of these pools could explain hypermetric scaling of heterophils in birds. A final possibility for scaling disparities between classes is how physical barriers scale with size. Bird feathers and mammalian fur both scale hypometrically [9], which might make large mammals and birds more susceptible to infection [6] and generally necessitate granulocyte hyperallometries. Perhaps the more negative *b* for skin exists because of its thinness and simplicity, which is distinct from mammalian skin [56].

We are far from resolving why birds require so much more heterophil-mediated protection at large body sizes, but we encourage additional work on the topic. It is also obscure whether the patterns we detected truly represent evidence for the Safety factor hypothesis, so we advocate that future work try to address this open question. Such insight will be useful for resolving how and why body size affects disease ecology and evolution. We expect that additional mechanistic studies will be useful, particularly the development of more precise tools for distinguishing lymphocyte classes or methods that capture better the functional immune variation (e.g., direct control of infections).

## Supporting information

Metadata

## Acknowledgements

We thank Samantha Oakey and Hannah Carey for help compiling data.

## Ethics

All data used in this study were obtained from the Species 360 database, which collects information from healthy, captive animals held at Association of Zoos and Aquariums (AZA) accredited facilities.

## Data, code and materials

Data used in this analysis will be available in Dryad digital repository.

## Author contributions

ECR, LBM and CJD conceived the idea, CJD oversaw collection of the data, ECR and CJD conducted the statistical analysis, and ECR, LBM and CJD wrote and revised the manuscript.

