## Supplementary tables and figures for "The impacts of body mass on immune cell concentrations in birds"

**Table S1.** Body mass and life history variables for each species included in our analysis (n=116). Sample size represents the number of samples taken, regardless of individual ID. Number of animals represents the unique number of individuals.

| *Scientific name* | *Common name* | *Sample size* | *Number animals* | *Maximum lifespan (yr)* | *Age at maturation (yr)* | *Interbirth interval (yr)* | *Avg clutch size* | *Incubation period* | *Maximal reproductive effort* | *Log(mass)* |
|---|---|---|---|---|---|---|---|---|---|---|
| *Tympanuchus cupido* | Greater prairie-chicken | 1493 | 934 | 5 | 1.0 | 1.0 | 13.0 | 23.0 | 0.013 | 2.937 |
| *Rhea americana* | Greater rhea | 141 | 105 | 13 | 2.0 | 1.0 | 30.0 | 25.0 | 0.014 | 4.362 |
| *Colinus virginianus* | Northern bobwhite | 125 | 95 | 6.4 | 1.0 | 1.0 | 15.0 | 23.0 | 0.015 | 2.250 |
| *Dendrocygna autumnalis* | Black-bellied whistling duck | 115 | 73 | 8.2 | 1.0 | 1.0 | 14.0 | 27.0 | 0.018 | 2.891 |
| *Callipepla gambelii* | Gambel's quail | 115 | 72 | 7.4 | 1.0 | 1.0 | 11.0 | 22.5 | 0.025 | 2.220 |
| *Netta rufina* | red-crested pochard | 124 | 95 | 8.8 | 1.0 | 1.0 | 10.0 | 27.0 | 0.028 | 3.048 |
| *Dromaius novaehollandiae* | Emu | 232 | 132 | 16.6 | 1.5 | 1.0 | 10.0 | 50.0 | 0.030 | 4.534 |
| *Dendrocygna bicolor* | Fulvous whistling duck | 286 | 158 | 11.2 | 1.0 | 1.0 | 13.0 | 25.0 | 0.030 | 2.879 |
| *Lophodytes cucullatus* | Hooded merganser | 448 | 306 | 15.7 | 2.0 | 1.0 | 11.0 | 31.0 | 0.039 | 2.790 |
| *Fratercula cirrhata* | Tufted puffin | 215 | 107 | 6 | 3.5 | 1.0 | 1.0 | 46.5 | 0.053 | 2.881 |
| *Bucephala islandica* | Barrow's goldeneye | 100 | 71 | 18 | 3.2 | 1.0 | 9.0 | 30.0 | 0.053 | 2.981 |
| *Bucephala clangula* | Common goldeneye | 60 | 46 | 18.4 | 4.0 | 1.0 | 8.7 | 30.0 | 0.053 | 2.962 |
| *Aix sponsa* | Wood duck | 295 | 195 | 22.5 | 1.0 | 2.0 | 12.0 | 31.0 | 0.054 | 2.313 |
| *Dendrocygna viduata* | White-faced whistling duck | 477 | 255 | 12 | NA | NA | 8.0 | 27.0 | 0.056 | 2.839 |
| *Athene cunicularia* | Burrowing owl | 339 | 168 | 11 | 0.8 | 1.0 | 6.0 | 29.0 | 0.056 | 2.281 |
| *Aythya americana* | Redhead | 216 | 144 | 22.6 | 1.0 | 1.0 | 13.0 | 26.0 | 0.062 | 3.032 |
| *Oxyura jamaicensis* | Ruddy duck | 408 | 285 | 13.6 | 1.0 | 1.0 | 8.0 | 24.5 | 0.062 | 2.784 |
| *Anas cyanoptera* | Cinnamon teal | 124 | 92 | 12.9 | 1.0 | 1.0 | 8.0 | 23.0 | 0.062 | 2.577 |
| *Bucephala albeola* | Bufflehead | 141 | 101 | 18.7 | 2.0 | 1.0 | 8.0 | 30.5 | 0.066 | 2.606 |
| *Anser canagicus* | Emperor goose | 44 | 40 | 12 | 3.5 | 1.0 | 5.0 | 24.5 | 0.067 | 3.447 |
| *Anas discors* | Blue-winged teal | 150 | 106 | 23.2 | 0.7 | 1.0 | 10.0 | 30.5 | 0.072 | 2.857 |
| *Aythya affinis* | Lesser scaup | 45 | 40 | 18.3 | 1.0 | 1.0 | 9.0 | 24.0 | 0.077 | 2.914 |
| *Anas clypeata* | Northern shoveler | 153 | 93 | 22.6 | 0.7 | 1.0 | 11.0 | 23.0 | 0.083 | 2.787 |
| *Myiopsitta monachus* | Monk parakeet | 47 | 32 | 22.1 | 2.0 | 1.0 | 7.0 | 31.0 | 0.090 | 2.079 |
| *Anas platyrhynchos* | Mallard | 296 | 161 | 29.1 | 1.0 | 1.0 | 11.0 | 27.0 | 0.091 | 3.034 |
| *Alopochen aegyptiaca* | Egyptian Goose | 190 | 99 | 25.5 | 2.0 | 1.0 | 8.5 | 29.0 | 0.092 | 3.273 |
| *Geococcyx californianus* | Greater roadrunner | 204 | 115 | 9 | 1.0 | 1.0 | 4.0 | 20.0 | 0.095 | 2.575 |

| | | | | | | | | | | |
|---|---|---|---|---|---|---|---|---|---|---|
| *Anser indicus* | Bar-headed goose | 167 | 104 | 20 | 3.0 | 1.0 | 5.5 | 29.0 | 0.103 | 3.347 |

| ***Scientific name*** | **Common name** | **Sample size** | **Number animals** | **Maximum lifespan (yr)** | **Age at maturation (yr)** | **Interbirth interval (yr)** | **Avg clutch size** | **Incubation period** | **Maximal reproductive effort** | **Log(mass)** |
|---|---|---|---|---|---|---|---|---|---|---|
| *Falco sparverius* | American kestrel | 224 | 97 | 17 | 1.0 | 1.0 | 5.0 | 30.0 | 0.103 | 2.063 |
| *Pygoscelis papua* | Gentoo penguin | 364 | 166 | 10.5 | 2.5 | 1.0 | 2.0 | 37.0 | 0.105 | 3.775 |
| *Aegolius acadicus* | Northern Saw-whet owl | 43 | 17 | 17.5 | 1.0 | 1.0 | 5.5 | 27.0 | 0.107 | 1.942 |
| *Cygnus melanocoryphus* | Black-necked swan | 402 | 181 | 20 | 2.0 | 1.0 | 4.6 | 35.0 | 0.109 | 3.679 |
| *Asio flammeus* | Short-eared owl | 47 | 14 | 21.8 | 1.0 | 1.0 | 5.8 | 29.0 | 0.120 | 2.540 |
| *Cygnus olor* | Mute swan | 128 | 58 | 40 | 3.0 | 1.0 | 5.0 | 60.0 | 0.121 | 4.031 |
| *Cygnus columbianus* | Tundra swan | 40 | 20 | 24.1 | 4.0 | 1.0 | 5.0 | 32.0 | 0.122 | 3.829 |
| *Butorides virescens* | Green heron | 65 | 29 | 11.6 | 1.0 | 1.0 | 4.0 | 20.0 | 0.126 | 2.326 |
| *Buteo lagopus* | Rough-legged buzzard | 52 | 13 | 18.8 | 2.5 | 1.0 | 4.0 | 31.0 | 0.127 | 2.980 |
| *Anas acuta* | Northern pintail | 287 | 190 | 27.4 | 1.0 | 2.0 | 8.0 | 23.0 | 0.132 | 2.976 |
| *Bubo scandiacus* | Snowy owl | 309 | 123 | 28 | 2.0 | 1.0 | 6.0 | 31.6 | 0.133 | 3.310 |
| *Cathartes aura* | Turkey vulture | 296 | 87 | 20.8 | 11.0 | 1.0 | 2.0 | 35.0 | 0.136 | 3.342 |
| *Aythya valisineria* | Canvasback | 129 | 81 | 29.5 | 1.0 | 1.0 | 8.0 | 25.0 | 0.137 | 3.080 |
| *Anser cygnoid* | Swan goose | 73 | 49 | 30 | 2.0 | 1.0 | 5.5 | 36.0 | 0.138 | 3.546 |
| *Pavo cristatus* | Indian peafowl | 388 | 226 | 23.2 | 3.0 | 1.0 | 5.0 | 28.0 | 0.139 | 3.623 |
| *Harpia harpyja* | Harpy eagle | 51 | 14 | 16.8 | 4.5 | 2.5 | 1.5 | 56.0 | 0.140 | 3.681 |
| *Anseranas semipalmata* | Magpie goose | 112 | 70 | 32 | 2.0 | 1.0 | 7.0 | 29.5 | 0.141 | 3.384 |
| *Parabuteo unicinctus* | Harris's hawk | 411 | 114 | 25 | 1.0 | 0.5 | 4.5 | 35.0 | 0.150 | 2.937 |
| *Cygnus buccinator* | Trumpeter swan | 470 | 304 | 32.5 | 5.5 | 1.0 | 5.0 | 34.5 | 0.152 | 4.045 |
| *Agapornis roseicollis* | Rosy-faced lovebird | 44 | 22 | 20 | 0.2 | 1.0 | 5.0 | 23.3 | 0.163 | 1.739 |
| *Megascops asio* | Eastern screech-owl | 429 | 182 | 20.7 | 1.0 | 1.0 | 4.0 | 29.0 | 0.164 | 2.215 |
| *Anser anser* | Greylag goose | 47 | 34 | 31 | 2.5 | 1.0 | 6.0 | 27.5 | 0.167 | 3.520 |
| *Spheniscus humboldti* | Humboldt penguin | 3031 | 606 | 17.5 | 3.0 | 0.5 | 2.0 | 41.0 | 0.175 | 3.641 |
| *Tyto alba* | Barn owl | 516 | 177 | 34 | 1.0 | 1.0 | 5.8 | 31.5 | 0.177 | 2.544 |
| *Polytelis alexandrae* | Alexandra's parrot | 65 | 44 | 23.9 | 4.0 | 1.0 | 5.0 | 21.0 | 0.181 | 2.017 |
| *Nycticorax nycticorax* | Black-crowned nightheron | 108 | 54 | 21.1 | 2.0 | 1.0 | 4.0 | 25.0 | 0.184 | 2.908 |
| *Buteo swainsonii* | Swainson's hawk | 84 | 26 | 19.5 | 2.0 | NA | 3.0 | 31.0 | 0.188 | 2.992 |
| *Corvus brachyrhynchos* | American crow | 116 | 36 | 20 | 2.0 | 1.0 | 5.0 | 18.0 | 0.189 | 2.704 |
| *Ciconia nigra* | Black stork | 77 | 16 | 31.3 | 4.0 | 1.0 | 4.0 | 35.0 | 0.190 | 3.477 |
| *Ciconia ciconia* | European white stork | 456 | 173 | 39 | 4.0 | 1.0 | 5.0 | 35.0 | 0.194 | 3.538 |

| Scientific name | Common name | Sample size | Number animals | Maximum lifespan (yr) | Age at maturation (yr) | Interbirth interval (yr) | Avg clutch size | Incubation period | Maximal reproductive effort | Log(mass) |
|---|---|---|---|---|---|---|---|---|---|---|
| *Falco peregrinus* | Peregrine falcon | 212 | 76 | 25 | 3.0 | 1.0 | 3.0 | 34.0 | 0.210 | 2.941 |
| *Melopsittacus undulatus* | budgerigar | 126 | 111 | 21 | 0.5 | 1.0 | 5.0 | 18.0 | 0.216 | 1.462 |
| *Egretta thula* | Snowy egret | 71 | 46 | 22.8 | 1.0 | 1.0 | 4.0 | 24.0 | 0.218 | 2.569 |
| *Aratinga solstitialis* | Sun parakeet | 235 | 149 | 22.5 | 2.0 | 1.0 | 3.5 | 25.0 | 0.225 | 2.047 |
| *Anser caerulescens* | Snow goose | 42 | 28 | 27.5 | 4.0 | 1.0 | 4.0 | 24.0 | 0.235 | 3.390 |
| *Ramphastos toco* | Toco toucan | 296 | 141 | 16.2 | 3.5 | 1.0 | 3.0 | 16.5 | 0.242 | 2.791 |
| *Coragyps atratus* | Black vulture | 112 | 38 | 25.5 | 8.0 | 1.0 | 2.0 | 35.0 | 0.243 | 3.334 |
| *Phalacrocorax auritus* | Double-crested cormorant | 40 | 30 | 22.5 | 2.0 | 1.0 | 3.0 | 26.5 | 0.248 | 3.259 |
| *Bubo virginianus* | Great horned owl | 438 | 133 | 29 | 2.0 | 1.0 | 3.0 | 33.5 | 0.261 | 3.161 |
| *Balearica regulorum* | Grey crowned crane | 949 | 349 | 27.2 | 3.0 | 1.0 | 3.0 | 29.0 | 0.269 | 3.544 |
| *Bubo bubo* | Eurasian eagle-owl | 182 | 51 | 68 | 2.0 | 1.0 | 3.0 | 77.5 | 0.280 | 3.429 |
| *Pulsatrix perspicillata* | Spectacled owl | 260 | 82 | 25 | 4.0 | 1.0 | 2.0 | 36.0 | 0.284 | 2.923 |
| *Spheniscus demersus* | Jackass penguin | 3049 | 858 | 27.3 | 4.0 | 1.0 | 2.0 | 40.0 | 0.284 | 3.496 |
| *Aegypius monachus* | cinereous vulture | 317 | 68 | 20 | 4.4 | 1.0 | 1.0 | 53.3 | 0.287 | 3.912 |
| *Buteo jamaicensis* | Red-tailed hawk | 426 | 153 | 30.7 | 3.0 | 1.0 | 3.0 | 30.0 | 0.298 | 3.052 |
| *Bubulcus ibis* | Cattle egret | 289 | 154 | 23 | 2.0 | 1.0 | 3.0 | 22.0 | 0.304 | 2.563 |
| *Eudyptula minor* | Little penguin | 306 | 190 | 25.6 | 3.0 | 1.0 | 2.0 | 35.5 | 0.310 | 3.061 |
| *Strix varia* | Barred owl | 173 | 72 | 24 | 2.0 | 1.0 | 2.3 | 30.5 | 0.310 | 2.855 |
| *Nymphicus hollandicus* | Cockatiel | 273 | 185 | 35 | 1.5 | 1.0 | 5.0 | 20.0 | 0.319 | 1.976 |
| *Somateria mollissima* | Common eider | 89 | 47 | 37.8 | 3.0 | 1.0 | 4.0 | 25.0 | 0.335 | 3.305 |
| *Cacatua alba* | White cockatoo | 102 | 40 | 26.9 | 5.5 | 1.0 | 2.0 | 30.0 | 0.345 | 2.756 |
| *Rhynchopsitta pachyrhyncha* | Thick-billed parrot | 798 | 200 | 30 | 2.0 | 1.0 | 3.0 | 26.0 | 0.346 | 2.512 |
| *Grus japonensis* | Red-crowned crane | 277 | 76 | 25.2 | 3.0 | 1.0 | 2.0 | 31.0 | 0.347 | 3.944 |
| *Cyanocitta cristata* | Blue jay | 50 | 22 | 26.2 | 1.0 | 1.0 | 4.0 | 17.0 | 0.350 | 1.944 |
| *Corvus albus* | White ibis | 68 | 20 | 27.6 | 3.0 | 1.0 | 3.0 | 22.0 | 0.357 | 2.719 |
| *Anthropoides virgo* | demoiselle crane | 228 | 106 | 27 | 6.0 | 1.0 | 2.0 | 28.0 | 0.362 | 3.383 |
| *Thraupis episcopus* | Blue-gray tanager | 206 | 148 | 9.5 | NA | NA | 2.0 | 13.0 | 0.365 | 1.544 |
| *Psittacus erithacus* | Grey parrot | 266 | 113 | 49.7 | 4.0 | 0.8 | 4.0 | 30.0 | 0.372 | 2.522 |
| *Geronticus eremita* | Northern bald ibis | 573 | 207 | 32.9 | 4.0 | 1.0 | 3.0 | 24.5 | 0.378 | 3.080 |
| *Ara macao* | Scarlet macaw | 582 | 212 | 33 | 3.5 | 1.5 | 3.0 | 24.5 | 0.378 | 3.006 |
| *Aptenodytes patagonicus* | King penguin | 222 | 88 | 26 | 5.0 | 1.0 | 1.0 | 54.0 | 0.382 | 4.070 |

| Scientific name | Common name | Sample size | Number animals | Maximum lifespan (yr) | Age at maturation (yr) | Interbirth interval (yr) | Avg clutch size | Incubation period | Maximal reproductive effort | Log(mass) |
|---|---|---|---|---|---|---|---|---|---|---|
| *Spheniscus magellanicus* | Magellanic penguin | 1290 | 257 | 30 | 3.4 | 1.0 | 1.5 | 41.0 | 0.422 | 3.615 |
| *Branta sandvicensis* | Hawaiian goose | 105 | 74 | 42 | 2.5 | 1.0 | 3.0 | 30.0 | 0.425 | 3.311 |
| *Fratercula corniculata* | Horned puffin | 106 | 46 | 20 | 4.0 | 1.0 | 1.0 | 35.0 | 0.444 | 2.726 |
| *Aquila chrysaetos* | Golden eagle | 355 | 114 | 48 | 5.5 | 1.0 | 2.0 | 42.0 | 0.494 | 3.630 |
| *Antigone canadensis* | Sandhill crane | 1958 | 419 | 36.6 | 4.5 | 1.0 | 2.0 | 30.0 | 0.518 | 3.623 |
| *Pelecanus occidentalis* | Brown pelican | 272 | 119 | 43 | 2.0 | 1.0 | 2.5 | 30.0 | 0.529 | 3.536 |
| *Corvus corax* | Common raven | 183 | 50 | 69 | 3.0 | 1.0 | 5.0 | 22.5 | 0.562 | 2.895 |
| *Ara ararauna* | Blue-and-yellow macaw | 599 | 225 | 43 | 3.5 | 1.5 | 2.5 | 26.0 | 0.575 | 3.051 |
| *Nestor notabilis* | Kea | 240 | 70 | 47 | 3.0 | 1.0 | 3.0 | 24.5 | 0.575 | 2.938 |
| *Grus americana* | Whooping crane | 1140 | 275 | 40 | 4.0 | 1.0 | 2.0 | 30.0 | 0.581 | 3.765 |
| *Amazona aestiva* | Blue-fronted parrot | 64 | 31 | 49 | 3.0 | 1.0 | 2.5 | 30.0 | 0.594 | 2.654 |
| *Haliaeetus leucocephalus* | Bald eagle | 1173 | 320 | 48 | 5.0 | 1.0 | 2.0 | 35.0 | 0.597 | 3.676 |
| *Threskiornis aethiopicus* | Sacred ibis | 307 | 151 | 37 | NA | 1.0 | 2.0 | 28.0 | 0.638 | 3.176 |
| *Ardeotis kori* | Kori bustard | 238 | 90 | 26 | 3.0 | 1.0 | 1.5 | 23.0 | 0.639 | 3.927 |
| *Amazona ochrocephala* | Yellow-crowned parrot | 50 | 13 | 56 | 3.0 | 1.0 | 3.0 | 25.5 | 0.667 | 2.643 |
| *Pelecanus onocrotalus* | Great white pelican | 127 | 56 | 51 | 3.5 | 1.0 | 2.0 | 32.5 | 0.709 | 3.979 |
| *Anodorhynchus hyacinthinus* | Hyacinth macaw | 449 | 157 | 38.8 | 8.0 | 1.0 | 1.5 | 26.5 | 0.747 | 3.124 |
| *Zenaida asiatica* | White-winged dove | 41 | 30 | 25 | 1.0 | 1.0 | 2.1 | 13.5 | 0.788 | 2.185 |
| *Pelecanus erythrorhynchos* | American white pelican | 354 | 163 | 54 | 3.0 | 1.0 | 2.0 | 30.0 | 0.823 | 3.752 |
| *Phoenicopterus ruber* | American flamingo | 2554 | 1020 | 30 | 4.0 | 1.0 | 1.0 | 29.5 | 0.852 | 3.491 |
| *Columba livia* | Rock dove | 157 | 125 | 35 | 0.4 | 1.0 | 2.0 | 18.5 | 0.888 | 2.550 |
| *Cacatua galerita* | Sulphur-crested cockatoo | 115 | 43 | 57 | 3.5 | 1.0 | 2.0 | 27.0 | 0.955 | 2.898 |
| *Phoenicopterus chilensis* | Chilean flamingo | 2311 | 990 | 36.7 | 6.0 | 1.0 | 1.0 | 29.5 | 1.007 | 3.357 |
| *Phoeniconaias minor* | Lesser flamingo | 381 | 203 | 41 | 3.5 | 6.5 | 1.0 | 29.5 | 1.042 | 3.176 |
| *Vultur gryphus* | Andean condor | 415 | 108 | 79 | 8.5 | 0.4 | 1.0 | 56.0 | 1.250 | 4.354 |
| *Phoenicopterus roseus* | Greater flamingo | 335 | 231 | 44 | 5.3 | 1.0 | 1.0 | 28.5 | 1.314 | 3.398 |
| *Probosciger aterrimus* | Palm cockatoo | 301 | 61 | 56.3 | 7.5 | 1.0 | 1.0 | 32.5 | 1.457 | 2.925 |

**Table S2.** Slope coefficients (b) and credible intervals (CI) of fixed effects in the omnibus model* for heterophil concentrations among 116 species of birds. Posterior mean is the mean of the posterior distribution.

| | **Posterior mean** | **l-95% CI** | **u-95% CI** | ***p*-value** |
|---|---|---|---|---|
| Intercept | 0.065 | -0.23 | 0.342 | 0.668 |
| Log(mass) | 0.210 | 0.122 | 0.297 | <0.001 * |
| Maximal reproductive effort | -0.375 | -1.089 | 0.457 | 0.338 |
| Maximum lifespan | 0.004 | -0.012 | 0.018 | 0.620 |
| Log(mass)*Maximal reproductive effort | 0.038 | -0.231 | 0.263 | 0.768 |
| Log(mass)*Maximum lifespan | -0.001 | -0.005 | 0.004 | 0.808 |

* $\beta_0 + \beta_1 \times \log_{10}(\text{Mass}) + \beta_2 \times \text{Max Repro} + \beta_3 \times \text{Max Longevity} + \beta_4 \times \log_{10}(\text{Mass}) \times \text{Max Repro} + \beta_5 \times \log_{10}(\text{Mass}) \times \text{Max Longevity}$

**Table S3.** Model predicting circulating leukocyte concentrations in birds and mammals. Model tested for the effects of body mass and class on $\log_{10}$-transformed lymphocyte and neutro-/heterophil concentrations.

| **Model** | **(adjusted) [95% CI]** | **(unadjusted) [95% CI]** | **Marginal $R^2$ [95% CI]** | **$R^2$** |
|---|---|---|---|---|
| Lymphocytes | | | | |
| $\beta_0 + \beta_1$ x $\log_{10}$(Mass) + Class + $\log_{10}$(Mass) x Class | 0.80 [0.65 : 0.91] | 0.72 [0.42 : 0.89] | 0.02 [$2.6e^{-3}$ : 0.646] | 0.80 |
| Heterophils | | | | |
| $\beta_0 + \beta_1$ x $\log_{10}$(Mass) + Class + $\log_{10}$(Mass) x Class | 0.91 [0.79 : 0.95] | 0.77 [0.43 : 0.89] | 0.11 [0.05 : 0.50] | 0.92 |

**Table S4.** Slope coefficients (b) and credible intervals (CI) of fixed effects in the direct bird/mammal analysis for lymphocytes and neutro-/heterophil concentrations among 116 species of birds and 259 species of mammals. Posterior mean is the mean of the posterior distribution.

| **Model** | **Posterior mean** | **l-95% CI** | **u-95% CI** | ***p*-value** |
|---|---|---|---|---|
| Lymphocytes | | | | |
| Intercept | 0.562 | 0.123 | 1.02 | 0.016* |
| Log(mass) | 0.01 | -0.039 | 0.061 | 0.686 |
| Class | 0.111 | -0.407 | 0.706 | 0.67 |
| Log(mass)*Class | -0.046 | -0.097 | 0.013 | 0.122 |
| Hetero-/Neutrophils | | | | |
| Intercept | 0.127 | -0.471 | 0.707 | 0.642 |
| Log(mass) | 0.194 | 0.141 | 0.243 | <0.001* |
| Class | 0.103 | -0.656 | 0.843 | 0.776 |
| Log(mass)*Class | -0.100 | -0.152 | -0.039 | <0.001* |

**Figure S1.** The body mass distribution of our 116 species of birds (panel A) compared to the body mass distribution of the world's bird species (6000 species; panel B). Mass distribution of extant bird species is used, with permission, from Blackburn and Gaston (1994). The Distribution of Body Sizes of the World's Bird Species. *Oikos* 70(1): 127-130.

A

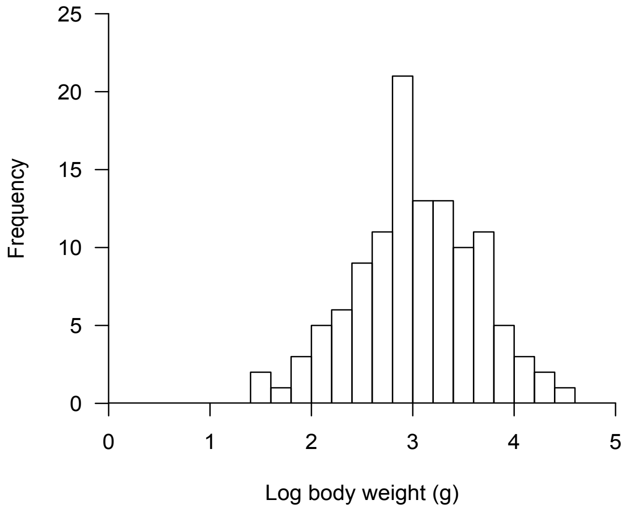


B

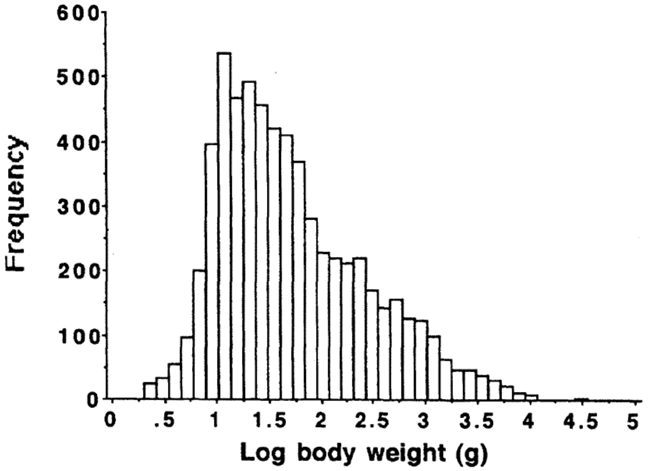

**Figure S2.** Time-rooted phylogenetic tree depicting evolutionary relationships among the avian species in this study. A total of 116 species were used for lymphocyte and heterophil models (all species, in red and black) whereas a subset of 88 species were used for the eosinophil concentrations (subset of species in red text). This tree was derived from Uyeda et al., 2017.

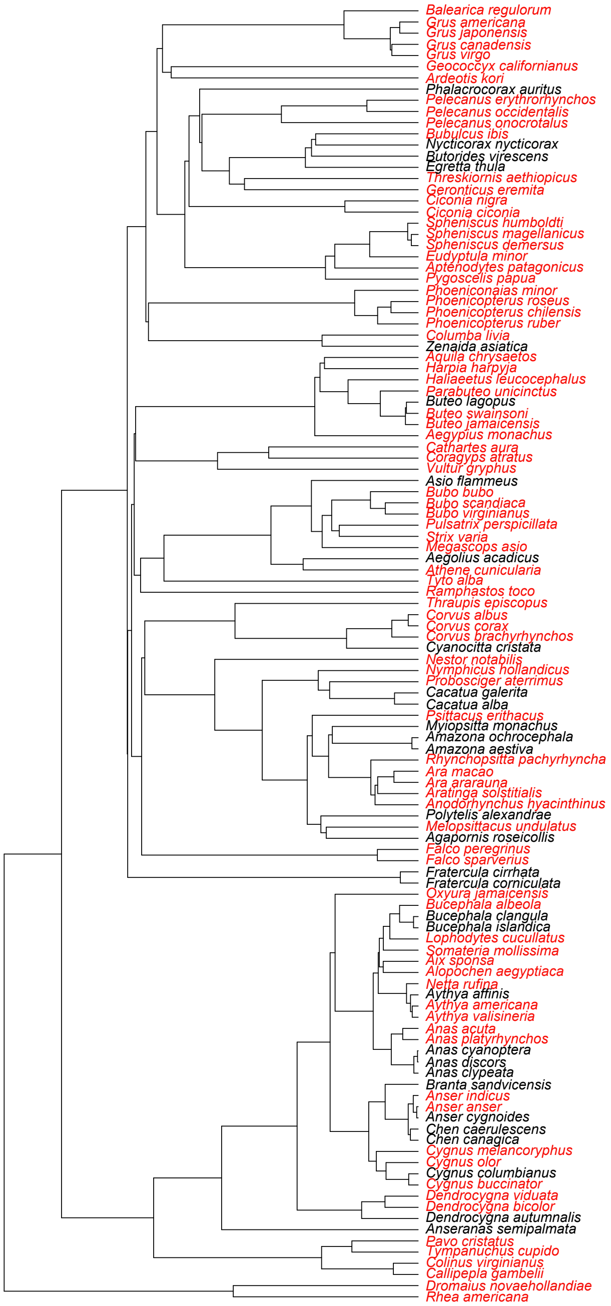

**Figure S3.** Correlations of $\log_{10}$-transformed body, maximal reproductive capacity, and maximum lifespan. The shortest-lived species is the Greater-prairie chicken (*Tympanuchus cupido*) and longest-lived the Andean condor (*Vultur gryphus*). $\text{Log}_{10}$-transformed body was positively correlated with maximal reproductive effort (S2a; t = 2.00, df = 137, p-value = 0.047), $\log_{10}$-transformed body was positively correlated with maximum lifespan (S2b; t = 3.24, df = 137, p-value = 0.001), and maximum lifespan was positively correlated with maximal reproductive effort (S2c; t = 12.31, df = 137, p-value < 0.001).

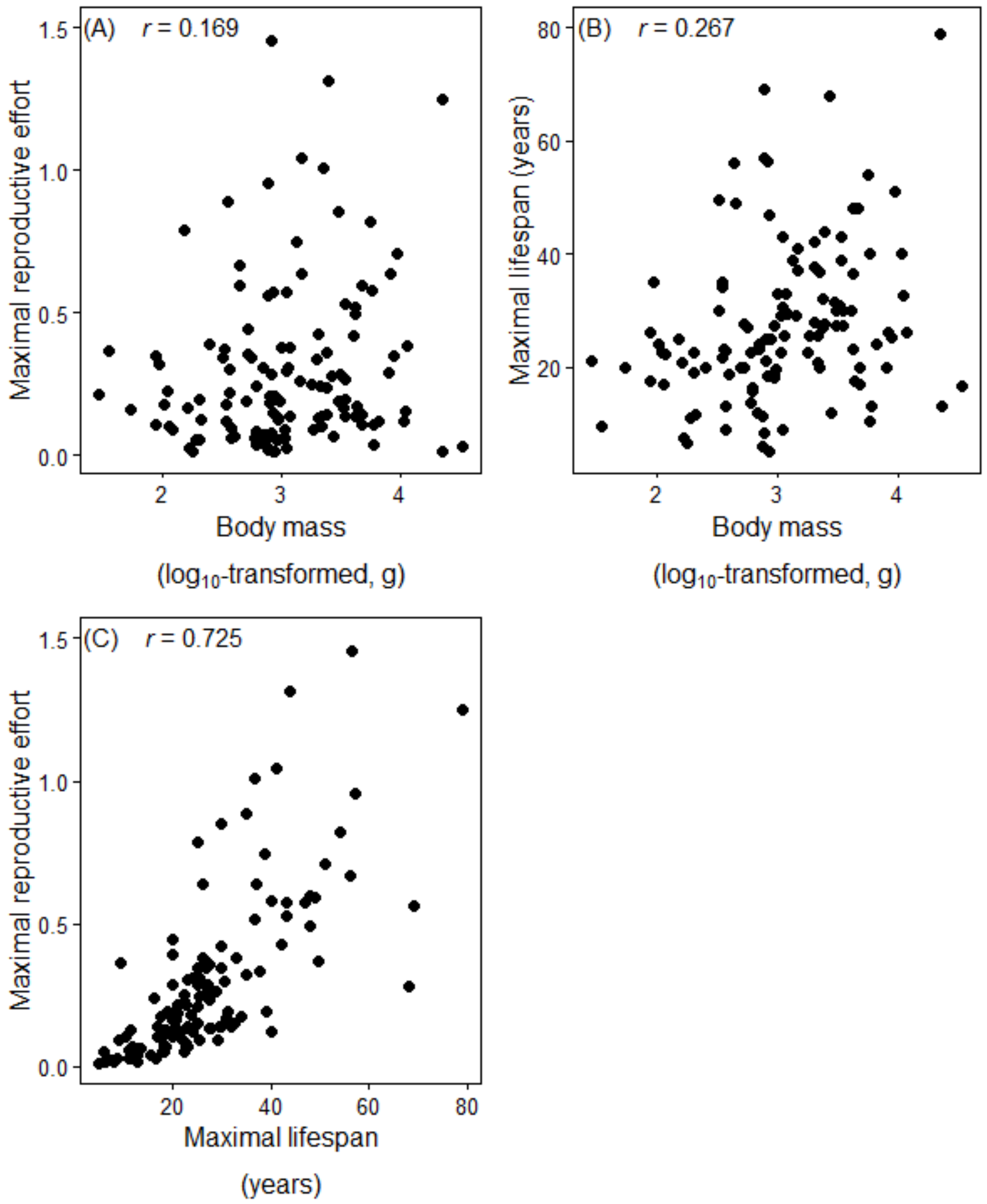

**Figure S4.** Observed scaling relationships between body mass and (A) lymphocytes and (B) eosinophils in birds. Equations: (a) $\log_{10}$(lymphocyte concentration) = 0.58±0.17 + 0.003±0.03($\log_{10}$body mass) and (b) $\log_{10}$(eosinophil concentration) = -0.76±0.32 + 0.09±0.1($\log_{10}$body mass). Shaded areas depict 95% credible intervals of the slope estimates from model 2 (mass-only). The intercept-only model was the best-performing model, hence these presented slopes are non-significant.

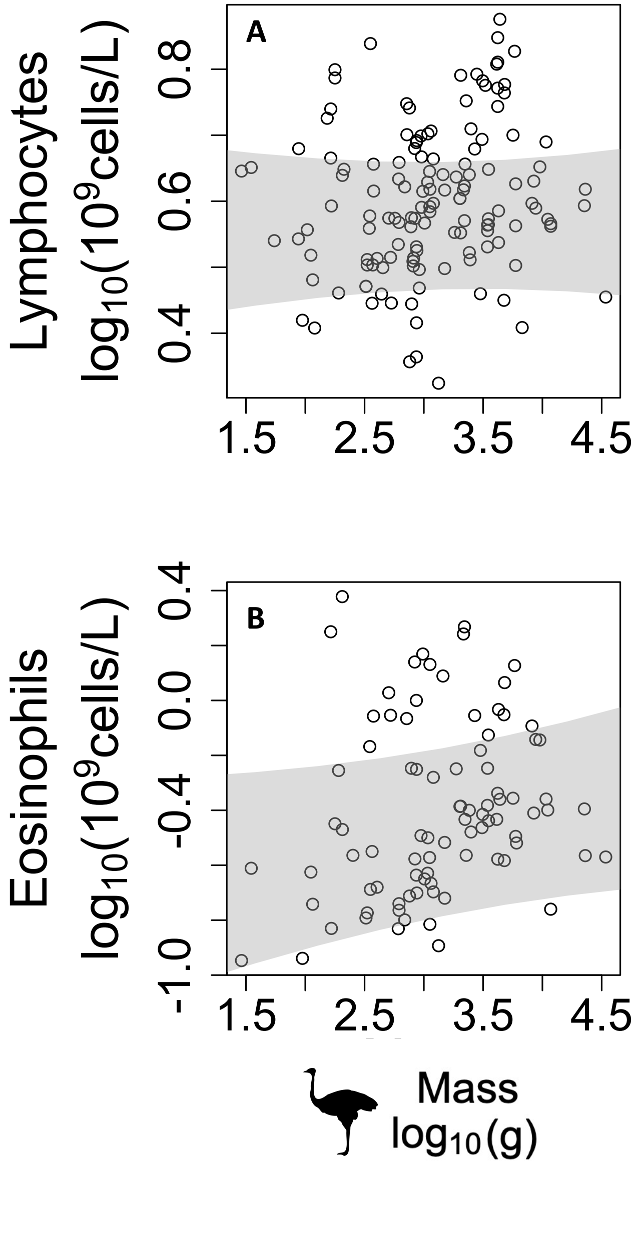
